# Neuronal and synaptic plasticity in the visual thalamus in mouse models of glaucoma

**DOI:** 10.1101/2020.10.23.352310

**Authors:** Matthew J. Van Hook, Corrine Monaco, Jennie C. Smith

## Abstract

Homeostatic plasticity plays important roles in regulating synaptic and intrinsic neuronal function to stabilize output following perturbations to circuit activity. In glaucoma, a neurodegenerative disease of the visual system commonly associated with elevated intraocular pressure (IOP), early disease is associated with altered synaptic inputs to retinal ganglion cells (RGCs), changes in RGC intrinsic excitability, and deficits in optic nerve transport and energy metabolism. These early functional changes can precede RGC degeneration and are likely to alter RGC outputs to their target structures in the brain and thereby trigger homeostatic changes in synaptic and neuronal properties in those brain regions. In this study, we sought to determine whether and how neuronal and synaptic function is altered in the dorsal lateral geniculate nucleus (dLGN), an important RGC projection target in the thalamus, and how functional changes relate to IOP. We accomplished this using patch-clamp recordings from thalamocortical (TC) relay neurons in the dLGN in two established mouse models of glaucoma – the DBA/2J (D2) genetic mouse model and an inducible glaucoma model with intracameral microbead injections to elevate IOP. We found that the intrinsic excitability of TC neurons was enhanced in D2 mice and these functional changes were mirrored in recordings of TC neurons from microbead-injected mice. Notably, many neuronal properties were correlated with IOP in older D2 mice, but not younger D2 mice or microbead-injected mice. The frequency of miniature excitatory synaptic currents (mEPSCs) was reduced in both ages of D2 mice, and vGlut2 staining of RGC synaptic terminals was reduced in an IOP-dependent manner in older D2 mice. Among D2 mice, functional changes observed in younger mice without elevated IOP were distinct from those observed in older mice with elevated IOP and RGC degeneration, suggesting that glaucoma-associated changes to neurons in the dLGN might represent a combination of stabilizing/homeostatic plasticity at earlier stages and pathological dysfunction at later stages.

## Introduction

Changes in synaptic and neuronal function are hallmarks of numerous neurological and neurodegenerative diseases (Wishart et al., 2006; Hall et al., 2015; Wang et al., 2016, 2017; Bae and Kim, 2017; Kim et al., 2017). In many cases, these are likely to be the result of early-stage homeostatic plasticity, which modulates neuronal response properties to maintain firing rate and compensate for altered circuit activity (Turrigiano, 2012; Wondolowski and Dickman, 2013). Such homeostatic modulation is accomplished by scaling presynaptic neurotransmitter release mechanisms, alterations in the balance of excitation and inhibition, modulation of post-synaptic neurotransmitter receptor complement and function, and tuning of intrinsic neuronal excitability and spiking behavior. In cases of disease or injury, homeostasis can make up for circuit dysfunction, but only to a point, after which disease processes are likely to overwhelm homeostasis and trigger unchecked dysfunction and degeneration (Orr et al., 2020). Understanding mechanisms of neuronal homeostasis in disease and injury is likely to provide insights into endogenous mechanisms with neuroprotective potential. Moreover, understanding pathological alterations in function is critical for shedding light on the timeline and mechanisms of neuronal loss and disease pathogenesis.

Glaucoma is a blinding neurodegenerative disease that strikes at retinal ganglion cells (RGCs), the output neurons of the retina (Calkins, 2012; Weinreb et al., 2014). It is commonly associated with elevated eye pressure, which damages RGC axons at the optic nerve head and culminates in the degeneration of their axon projections to visual regions of the brain and apoptotic loss of cell bodies in the retina. However, glaucoma-associated changes to visual function are likely the result of more than RGC and axonal degeneration, as cell loss is preceded by numerous changes in the structure and function of RGC-associated circuits. Within the retina, for instance, pressure elevation upregulates Na^+^ channel expression and spiking behavior of RGCs, alters the size and complexity of their dendritic fields, changes receptive field properties, leads to changes in post-synaptic receptor composition, and alters the responses of upstream retinal circuits (Della Santina et al., 2013, 2013; Frankfort et al., 2013; Wang et al., 2014; Pang et al., 2015; Ou et al., 2016; Bhandari et al., 2019; McGrady et al., 2020; Sladek and Nawy, 2020). Within the optic projection and at RGC output sites in the brain, eye pressure and glaucomatous injury lead to alterations in optic nerve glial function and energy demands (Baltan et al., 2010; Inman and Harun-Or-Rashid, 2017; Jassim et al., 2019; Cooper et al., 2020), RGC axon terminal swelling and atrophy (Smith et al., 2016), changes in axon terminal mitochondrial health (Smith et al., 2016), altered synaptic vesicle release properties (Bhandari et al., 2019), and post-synaptic neuronal dendritic remodeling and somatic atrophy (Gupta and Yücel, 2003; Gupta et al., 2007, 2009, Liu et al., 2011, 2014; Bhandari et al., 2019). It is likely that such changes represent a mix of both homeostatic attempts at preserving retinal output fidelity and pathological alterations in neural function in glaucoma. Overall, the relative timing, balance, and interplay of these two possible processes remains unknown.

The goal of this study was to test the hypothesis that glaucoma progression triggers changes in neuronal and synaptic function in the dorsal lateral geniculate nucleus (dLGN), a major RGC projection target in the thalamus that underlies conscious, image-forming vision by receiving and processing RGC signals and relaying them to the primary visual cortex (Kerschensteiner and Guido, 2017; Seabrook et al., 2017). Long considered a simple relay structure, recent evidence has highlighted how changes in visual activity can alter dLGN synapses and neuronal function (Rose and Bonhoeffer, 2018). For instance, the maturation of neuronal excitability, dendritic structure, and synaptic transmission is regulated by visual experience and retinal inputs during development and young adulthood (Hooks and Chen, 2006; Hong and Chen, 2011; Seabrook et al., 2013; Louros et al., 2014; El-Danaf et al., 2015; Liang and Chen, 2020). Moreover, several studies have documented examples of functional alterations occurring in dLGN neurons and synapses in response to altered sensory input and in cases of disease and injury (Krahe and Guido, 2011; Araújo et al., 2017; Sommeijer et al., 2017; Rose and Bonhoeffer, 2018; Bhandari et al., 2019). This raises the possibility that RGC injury and dysfunction occurring in glaucoma can trigger compensatory homeostatic plasticity in dLGN synapses and neurons.

Therefore, we set out to determine whether and how changes to neuronal function relate to eye pressure and glaucomatous RGC degeneration to shed light on the links between eye pressure, neuronal homeostasis, and dysfunction in glaucoma. This was accomplished by using two complementary mouse models of glaucoma and probing for structural and functional alterations to neurons and synapses in the dLGN. Ultimately, we find changes in excitatory synaptic transmission onto the principal dLGN relay neurons. Additionally, glaucoma enhances their excitability in a manner associated with eye pressure at a later phase of the disease. At a later time point, we find RGC synaptic terminal loss associated with IOP and atrophy of neuronal cell bodies in the dLGN, which are likely to be signs of later-stage disease pathology. This implies that the dLGN in glaucoma is characterized by a potentially homeostatic modulation of function occurring early in the disease process, followed by later alterations and degeneration that are linked to disease severity.

## Materials and Methods

### Animals

Animal protocols were approved by the Institutional Animal Care and Use Committee at the University of Nebraska Medical Center. DBA/2J (D2; Jax# 000671) were used as an inherited model of glaucoma (John et al., 1998; Libby et al., 2005; Howell et al., 2007) and bred in-house or purchased from Jackson Labs. A strain-matched control line that contains a wild allele of the *Gpnmb* gene and does not develop elevated eye pressure or glaucoma was used as a control (DBA/2J^*Gpnmb+*/SjJ^; D2-control; Jax# 007048) (Howell et al., 2007). For microbead injection experiments, we used mice produced as a cross of Opn4^Cre/Cre^ (Ecker et al., 2010) and Ai32 (Jax #024109) (Madisen et al., 2012) lines (Opn4-Cre;Ai32) (Bhandari et al., 2019). Mouse were housed in a 12/12 hour light/dark cycle and provided with food and water *ad libitum*.

### Microbead occlusion model

To induce ocular hypertension, fluorescently-tagged polystyrene microspheres (10 micron, Invitrogen F8836) were bilaterally injected into the anterior chambers of Opn4^Cre^;Ai32 mice at ~6-8 weeks of age (Sappington et al., 2010; Calkins et al., 2018; Bhandari et al., 2019). This procedure was performed under isoflurane anesthesia and following instillation of anesthetic eye drops (0.5% proparacaine, Akorn, Lake Forest, IL). Pupils were dilated with 1% tropicamide eye drops. In total a small volume of beads (~1-2 μL) at a concentration of ~14×10^6^ beads/mL was injected using a glass micropipette.

### Intraocular pressure measurements

Intraocular pressure (IOP) was monitored using a Tonolab rebound tonometer (iCare, Vantaa, Finland) in mice lightly anesthetized with isoflurane, as we have described previously (Bhandari et al., 2019). IOP was measured approximately monthly in D2 and D2-control mice beginning around 2-4 months of age and was measured prior to and at 2 days, 1, 2, and 4 weeks post-injection for microbead-injected mice. For assessing the cumulative injury effects of IOP in D2 mice, we calculated a “3-month cumulative IOP”, which for a given mouse, is the sum of monthly IOP measurements taken over a three month-span (IOP measurements at 7, 8 and 9 months of age) and averaged across the two eyes. For microbead injected mice, we calculated a cumulative ΔIOP over baseline for each mouse as the average across the two eyes of 1 week, 2, week, and 4-week post-injection IOP measurements over the pre-injection baseline.

### Brain slice patch-clamp electrophysiology

250-micron thick coronal brain slices containing the dLGN were acutely prepared using the “protected recovery” method (Ting et al., 2014, 2018), as we have described previously (Bhandari et al., 2019; Van Hook, 2020). Following euthanasia by CO_2_ asphyxiation and cervical dislocation, brains were submerged in a slush of artificial cerebrospinal fluid (aCSF) comprised of (in mM) 128 NaCl, 2.5 KCl, 1.25 NaH_2_PO_4_, 24 NaHCO_3_, 12.5 glucose, 2 CaCl_2_, and 2 MgSO_4_ and continuously bubbled with a mixture of 5% CO_2_ and 95% O_2_. Slices were prepared on a vibrating microtome (Leica VT1000S) and hemisected through the midline. Slices were then incubated for ~12 minutes at 33°C in an N-methyl-D-glucamine-based solution (in mM: 92 NMDG, 2.5 KCl, 1.25 NaH_2_PO_4_, 25 glucose, 30 NaHCO_3_, 20 HEPES, 0.5 CaCl_2_, 10 MgSO_4_, 2 thiourea, 5 L-ascorbic acid, and 3 Na-pyruvate), after which they were transferred to a solution of room-temperature aCSF and allowed to recover for >1 hour prior to beginning recording.

dLGN slices were transferred to a recording chamber on a fixed-stage upright microscope (Olympus BX51-WI) and superfused with aCSF at a rate of 2-4 mL/min. For recordings from D2 and D2-control mice, the aCSF was warmed to 30-33°C using an inline heater. For recordings from microbead-injected mice, experiments were performed at room temperature (~23°C) and the aCSF was supplemented with 60 microM picrotoxin. Thalamocortical relay (TC) neurons located in the dLGN core (>100 microns from the dorso-lateral surface of the dLGN) were targeted for whole-cell recording based on soma size and shape and distinguished from interneurons by the presence of a pronounced low-voltage activated T-type Ca^2+^ current. For voltage-clamp experiments, the patch pipette solution was comprised of (in mM) 120 Cs-methanesulfonate, 2 EGTA, 10 HEPES, 8 TEA-Cl, 5 ATP-Mg, 0.5 GTP-Na_2_, 5 phosphocreatine-Na_2_, 2 QX-314 (pH = 7.4, 275 mOsm) while for current-clamp experiments, the pipette solution was comprised of (in mM) 120 K-gluconate, 8 KCl, 2 EGTA, 10 HEPES, 5 ATP-Mg, 0.5 GTP-Na_2_, 5 phosphocreatine (pH = 7.4, 275 mOsm). Miniature excitatory post-synaptic currents (mEPSCs) were recorded in the absence of stimulation at a holding potential of −70 mV. TC neuron spiking was evoked using a series of depolarizing current injections (+40 to +560 pA, 500 ms) while membrane properties were monitored using hyperpolarizing injections (−20 to −100, 500 ms).

### Electrophysiology analysis

Action potentials from current-clamp experiments were detected and counted using the “event detection” function of Clampfit. Input resistance (Rin) was measured as the slope of a straight line fit to the voltage deflection amplitudes evoked by current stimuli of −20 and −40 pA. Membrane time constant (τ_m_) was measured with a single exponential fit to the voltage deflection evoked by a −20 pA step and was used along with Rin to calculate the membrane capacitance (Cm = τ_m_ /Rin). mEPSCs were detected and analyzed using MiniAnalysis software (Synaptosoft, Fort Lee, NJ). For each cell, the first ~100 detected events were analyzed. Using the same number of events for each recorded cell avoids biasing the cumulative distribution analysis of event amplitude and inter-event intervals. All reported voltages are corrected for a measured −14 mV liquid junction potential for the K-based pipette solution and a −10 mV liquid junction potential for the Cs-based solution.

### Immunofluorescence staining

For labeling RGC somata, we performed immunofluorescence staining of flat-mount retinas with a guinea pig-anti-RBPMS antibody (PhosphoSolutions, 1832-RBPMS, 1:500, RRID: AB_2492226) (Rodriguez et al., 2014). Position in the retina was determined by staining with a primary antibody sensitive to S-opsin (rabbit-anti-s-opsin, 1:500, Millipore ABN1660) (Sondereker et al., 2018; Stabio et al., 2018), which is expressed in a gradient along the dorsal-ventral axis of the retina (Applebury et al., 2000). Retinas were dissected into oxygenated aCSF or Ames solution. Four relieving cuts were made, and the retina was mounted on a nitrocellulose membrane (type AAWB, 0.8 micron pore size, Millipore, Burlington, MA, USA) and fixed by immersion in 4% paraformaldehyde for 30 mins. Retinas were washed, blocked and permeabilized in a solution of 1% triton X-100, 0.5% DMSO, 5.5% donkey serum, and 5.5% goat serum for 1 hour before being incubated overnight at 4°C in the same solution plus addition of the primary antibodies. Retinas were then washed 6x, blocked/permeabilized again, and incubated with an AlexaFluor 568-conjugated goat-anti-guinea pig and an AlexaFluor 488-conjugated donkey-anti-rabbit secondary antibodies for 2 hours at room temperature. After washing 3x in PBS, retinas were removed from the nitrocellulose membranes, mounted on Superfrost Plus slides, and coverslipped with VectaShield Hardset.

For immunofluorescence staining of dLGN sections, mice were euthanized and brains dissected, as described above. After a brief rinse in PBS, they were immersed in 4% paraformaldehyde for 4 hours after which they were rinsed 3x in PBS and cryoprotected for 1-3 nights by immersion in 30% sucrose in PBS at 4°C. Brains were embedded in 3% agar, cut into 50 micron sections with a Leica VT1000S vibratome, mounted on SuperFrost Plus slides and stored at −20°C. After blocking/permeabilization in PBS containing 0.5% TritonX-100, 5.5% (donkey and/or goat serum), slices were stained using either a rabbit-anti-vGlut2 primary antibody (1:250, Cedarlane/Synaptic Systems #135403, RRID: AB_887883) or a combination of guinea pig-anti-NeuN polyclonal antibody (Millipore ABN90, RRID:AB_11205592, 1:500) and rabbit-anti-GAD65/67 polyclonal antibody (1:500, Millipore G5163, RRID: AB_477019) overnight at 4°C in a humidified chamber. For staining with the guinea pig primary antibody, we used a blocking solution containing both goat and donkey serum. Secondary antibodies were goat- or donkey-raised AlexaFluor 488 or 568-conjugated antibodies and were used at a concentration of 1:200. Slides were then washed 3x in PBS followed by 1x in dH2O and coverslipped using VectaShield Hardset.

### Imaging and image analysis

Imaging was performed with a 2-photon microscope with a 20x water-immersion objective (Scientifica) with the laser tuned to 800 nm. vGlut2 images were acquired in frames of 185×185 μm (5.54 pixels/μm) with 0.5 μm z-axis spacing from the dLGN core. Four images per focal plane were averaged and a maximum intensity projection was created from five planes to give an effective z-slice thickness of 2.5 microns. The vGlut2 signal was automatically thresholded and puncta were detected using the Synapse Counter plug-in in ImageJ (Dzyubenko et al., 2016) with a size threshold of 9 μm^2^.

For imaging RGCs, RBPMS-stained retinas were imaged in frames of 350 x 350 μm (1.04 pixels/μm). A series of images were acquired through the ganglion cell layer at 1 micron spacing in four quadrants of the central and peripheral retina (~500 and 1700 μm from the optic nerve head, respectively). Each of the four quadrants were identified as temporal, nasal, ventral, or dorsal based on the dorsal-ventral gradient of s-opsin cone labeling (Applebury et al., 2000). Four images per focal plane were averaged for analysis and RGCs were counted using the Cell Counter plug-in in ImageJ.

NeuN and GAD65/67-stained neurons in the dLGN core were imaged in frames of 185 x 185 μm (5.54 pixels/μm) with 1 micron z-axis spacing. Four images per plane were acquired and averaged for analysis. To measure cross-sectional soma area, NeuN-stained cell bodies were traced by hand in the image plane of each cells’ greatest area. After tracing, the regions of interest were superimposed on the GAD65/67 signal and GAD65/67-positive cell bodies were excluded from soma area analysis.

### Statistical analysis

Data are presented as mean + SEM unless indicated otherwise. Statistical significance was assessed using several different approaches. A two-tailed nested t-test was performed using GraphPad Prism 8 for electrophysiology experiments in which multiple cells were recorded from multiple animals, as indicated below. Otherwise, an unpaired two-tailed Student’s t-test was used, as indicated. Significance threshold was set at p<0.05 for t-tests. Differences in cumulative distributions of mEPSC inter-event intervals, mEPSC amplitudes, vGlut2 punctum sizes, and dLGN TC neuron soma sizes were assessed using a Kolmogorov-Smirnov (K-S) test in ClampFit 10. For K-S tests, p<0.0005 was considered statistically significant. A linear regression was used to test for correlations of mEPSC, excitability, RGC density, and vGlut2 staining with intraocular pressure. For correlation analysis, the mean values of each parameter for each mouse was used for the fit. A p-value <0.05 was considered a statistically significant correlation.

## Results

### dLGN TC neuron excitability is enhanced in D2 mice

To determine whether glaucoma and IOP affect the function of neurons in the dLGN, we made use of the DBA/2J (D2) line of mice. These mice displayed elevated IOP beginning around 7 months of age when compared to a strain-matched control mouse line (DBA/2J^*Gpnmb+/SjJ*^; D2-control; Figure 1). The IOP elevation was variable in D2 mice and peak IOP within the population (aged 9-10 months) ranged from 11 mmHg to 39 mmHg (n = 52 eyes, 26 mice). This distribution was significantly different than the maximum IOP values from D2-control mice (p<0.00001, K-S test). At 9 months of age, for instance, IOP was 20.2 + 6.2 mmHg (Mean + SD; n = 32 eyes, 16 mice) for D2 compared to 11.9 + 2.4 for 9 month-old D2-control mice (n = 16 eyes, 8 mice; p < 0.00001). This timeline and variability in IOP is consistent with prior studies of D2 mice (John et al., 1998; Libby et al., 2005; Inman et al., 2006).

**Figure 1.**
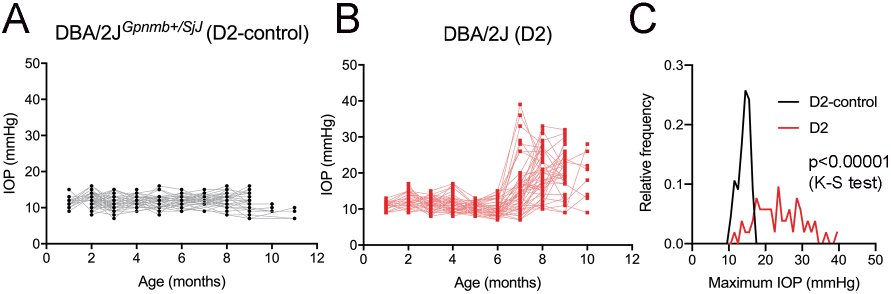
Eye pressure is elevated in DBA/2J mice. A&B) Intraocular pressure (IOP) measurements from DBA/2J^*Gpnmb+/SjJ*^ (D2-control, A) mice (n = 62 eyes from 31 mice) and DBA/2J (D2, B) mice (n = 66 eyes from 33 mice). C) Histogram of peak IOP values obtained from D2 and D2-control mice. D2-control IOPs were narrowly distributed (median = 14 mmHg) while the peak IOPs from D2 mice showed a broader distribution (median = 23 mmHg).

Glaucoma is triggered by injury to retinal ganglion cell axons, which comprise the optic nerve and are responsible for carrying information to visual centers of the brain. Because of this, we next sought to determine whether D2 mice display changes in the function of neurons in the dLGN, a key RGC projection target for conscious vision, and relate those changes to IOP. To do this, we targeted thalamocortical (TC) relay neurons in the dLGN for whole cell current clamp recording in acute coronal dLGN brain slices (Figure 2). Experiments were performed in slices from D2 and D2-control mice at four and nine months (4m and 9m) of age in order to compare function at an earlier (pre-ocular hypertension/pre-OHT) and a later (OHT) time point.

**Figure 2.**
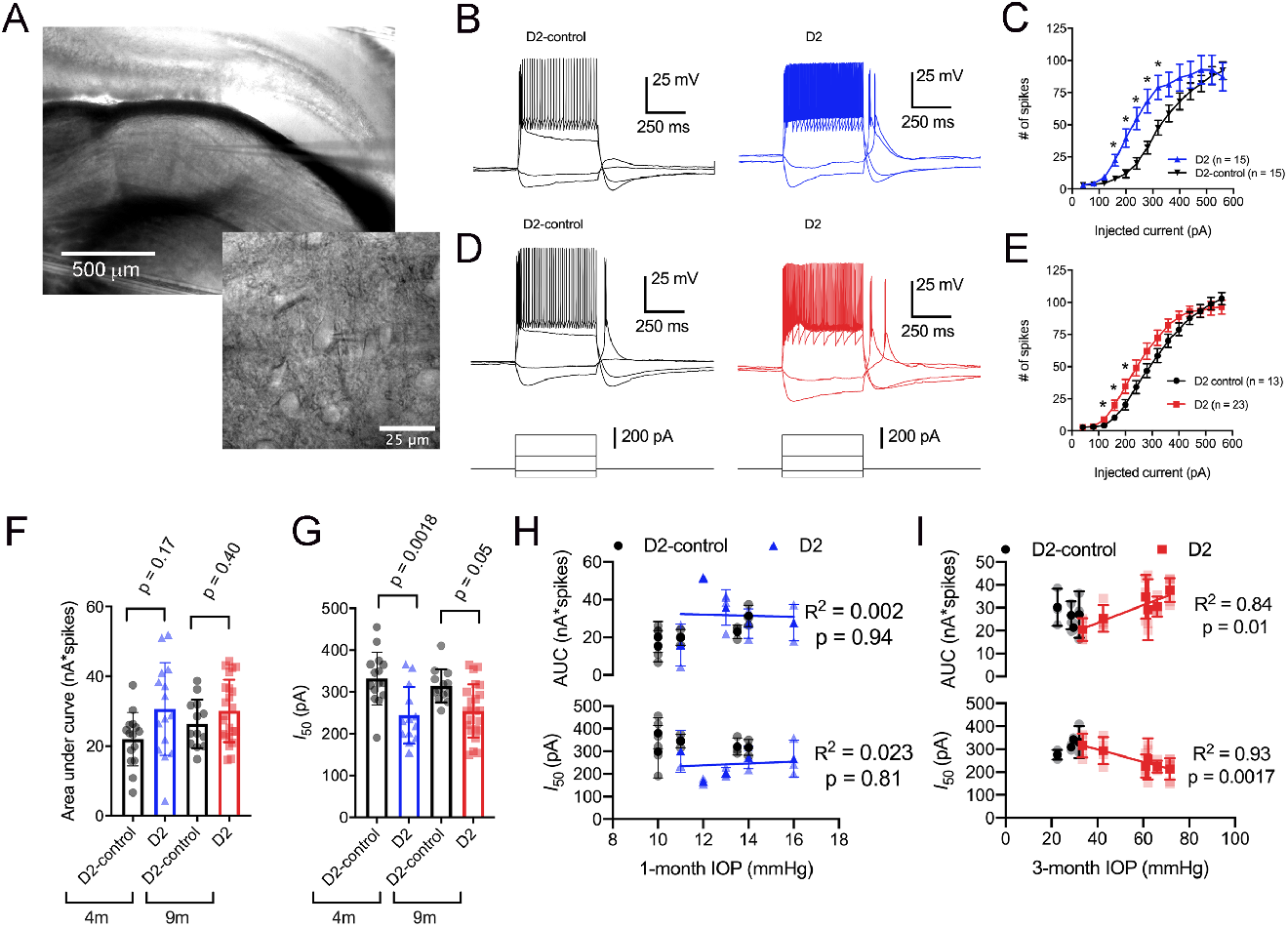
Enhancement of TC neuron spiking in D2 mice. A) Left, microscopy image of dLGN with patch-clamp electrode in coronal brain slice. Right, higher magnification image showing a TC neuron with patch clamp electrode. B) Evoked spiking responses to depolarizing current injection (+120 and + 320 pA) and voltage deflections to hyperpolarizing current injections (−20 and −80 pA) in 4m D2 mice (15 cells, 5 mice) and D2-control mice (n = 15 cells, 5 mice). C) Plot of current injection and counts of evoked spikes (mean + SEM) show that TC neurons from 4m D2 mice fired more spikes (*p<0.05, unpaired t-test). D) Spiking and hyperpolarizing voltage responses from 9m D2-control and D2 mice. E) Similar to panel C, TC neurons from 9m D2 mice (n = 23 cells, 6 mice) were slightly more excitable than from D2-control mice (n = 13 cells, 4 mice). F) Mean + SD of the area under the curve of spiking responses and data points from individual cells from 4m and 9m D2 and D2-control mice. Significance was assessed using a nested t-test approach. G) Mean + SD and individual cell data points of the half-maximal current injection from a Boltzmann fit to the current-spiking relationships. H) There was no significant correlation of AUC or I_50_ with IOP (measured at 4m of age) as assessed with a linear regression for 4m D2 mice. Regression was performed on mean values of AUC and I_50_ for each mouse rather than on individual cells. I) In 9m D2 mice, there was a significant correlation of AUC and I_50_ with a three-month cumulative IOP averaged across two eyes for each mouse.

When we used depolarizing current injections (500 ms, 40-560 pA; Figure 2B-E), we found that spiking was enhanced in 4-month old (4m) D2 mice compared to D2-controls. For instance, a 240 pA current injection evoked 19 + 4 action potentials in D2-control mice (n = 15 cells, 5 mice) whereas TC neurons from age-matched D2 mice fired 55 + 9 action potentials (n = 15 cells, 5 mice; p = 0.0016). We further quantified the enhanced excitability by measuring the area under the curve (AUC) of the current-spike relationship as well as the half-maximal current stimulus of a Boltzmann fit (I_50_) (Figure 2F&G). In this analysis, the AUC from 4m D2 cells was elevated (31 + 3 nA*spikes) relative to D2 controls (22 + 2 nA*spikes), although the difference was not significant (p = 0.17). The I_50_ was significantly shifted left, from 332 + 16 pA for D2-controls to 244 + 17 pA for D2 mice (p=0.0018).

When we performed similar experiments with 9m D2 and D2-control mice, the difference in excitability was similar although differences generally did not reach statistical significance. For instance, the AUC was 26 + 2 nA*spikes in 9m D2-control mice (n = 13 cells, 4 mice) and 30 + 2 nA*spikes in 9m D2 mice (n = 23 cells, 6 mice; p = 0.4). Additionally, the I_50_ was 314 + 11 pA in D2-controls and 254 + 13 in D2’s (p = 0.048).

Notably, we did find that there was considerable variability of the spiking in 9m D2 TC neurons relative to D2-controls. For instance, the SD of the I_50_ was 64 pA for 9m D2 and 39 pA for D2-control while for AUC, the SD was 9.0 for D2 and 6.9 for D2-control. As shown above, there was also wide variability in D2 mouse IOP values at 9 months. We turned this to our advantage by testing whether AUC and I_50_ were related to the cumulative IOP experienced by the D2 mouse visual system by plotting AUC and I_50_ against the 3-month cumulative IOP (Figure 2H&I). When we did this, we found that the AUC positively correlated with cumulative IOP (p = 0.01) while the I_50_ negatively correlated with cumulative IOP (p = 0.002). In contrast, IOP was not elevated in the 4-month old D2 mice and there was no significant correlation of AUC or I_50_ with IOP (p>0.05). Thus, higher eye pressure in aged D2 mice was associated with greater dLGN TC neuron excitability.

We next sought to probe the cellular mechanisms underlying this change in neuronal excitability (Figure 3) and found that the increase in TC neuron excitability in both 4m and 9m D2 mice was associated with changes in membrane properties that support increased action potential firing, although the effects were slightly different at each age. At 4m, D2 TC neurons were slightly depolarized relative to D2-controls (D2: −78.2 + 0.7 mV; D2-control: −81.3 + 0.6 mV; p = 0.011), which brings their membrane potential closer to action potential threshold. Additionally, input resistance (Rin), measured with a linear fit to voltage responses to hyperpolarizing current injections (Figure 3A&B), was higher in 4m D2 mice (D2: 288 + 20 MW; D2-control: 221 + 17 MΩ; p=0.018). Membrane capacitance (Cm), an electrical measure of cell surface area, was not significantly different between D2 and D2-control at 4 months (D2: 105.2 + 7.1 pF; D2-control: 117.3 + 8.0 pF; p = 0.27).

**Figure 3.**
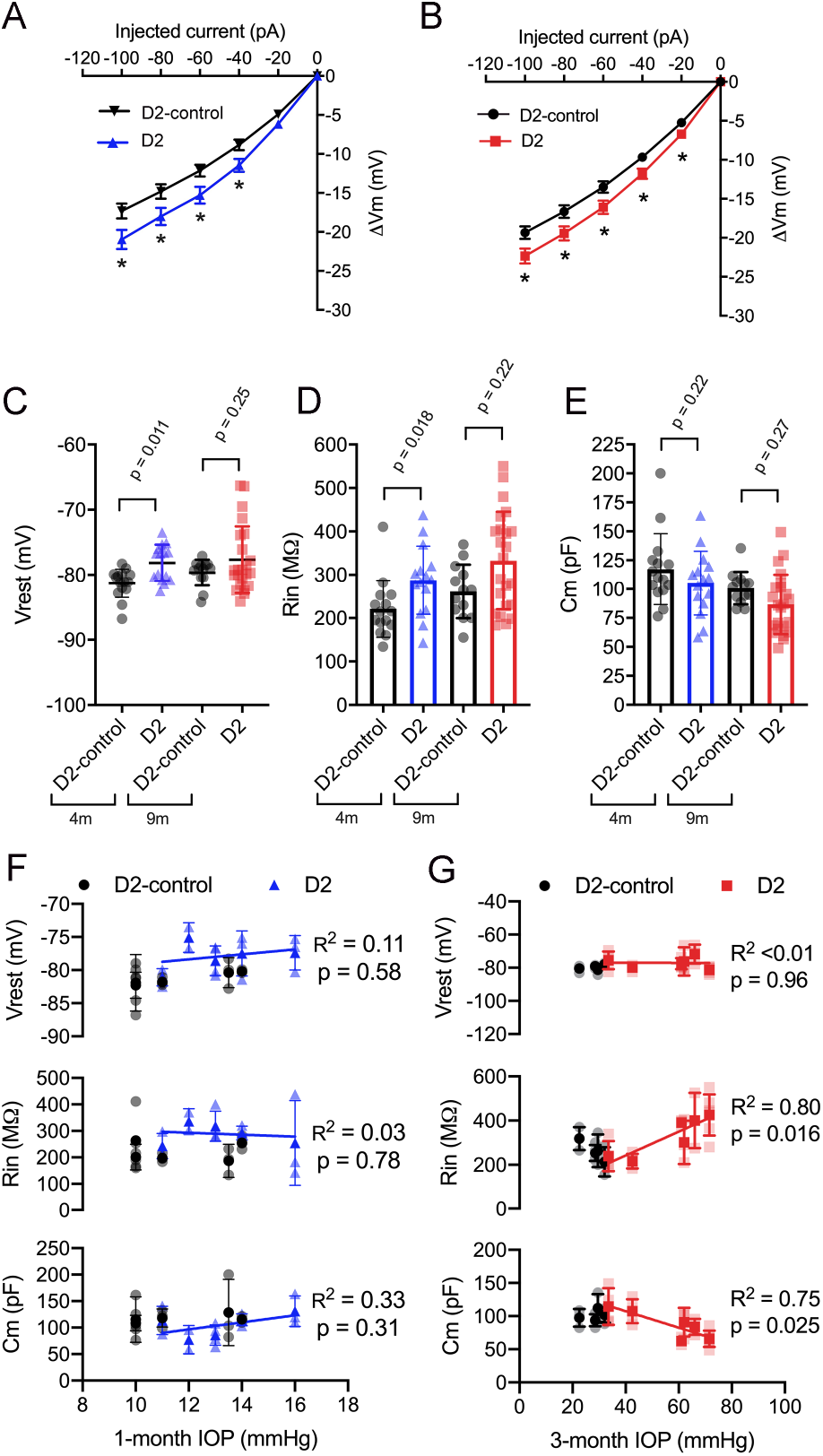
Changes to passive membrane properties support enhanced TC neuron excitability in D2 mice. A&B) Plots of voltage deflection evoked by hyperpolarizing current injections in 4m (A) and 9m (B) D2-control and D2 mice. In D2 mice, hyperpolarizing current injections evoked larger voltage changes (*p<0.05, unpaired t-test). C) Resting membrane potential (Vrest) measured in D2 and D2-control mice at 9m and 4m. Vrest was significantly depolarized in 4m D2 mice, but not 9m D2 mice when compared with age-matched controls (nested t-test). Input resistance (Rin), measured using a linear fit to voltage deflections evoked by −20 and −40 pA, was significantly elevated in 4m D2 TC neurons, but not for 9m when compared using a nested t-test. E) Membrane capacitance was not significantly different between D2-control and D2 mice at 4m or 9m (nested t-test). F) There was no significant correlation of Vrest, Rin, or Cm with IOP at 4m as assessed using a linear regression. G) Although there was no significant correlation of Vrest in 9m D2 TC neurons with the 3-month cumulative IOP, Rin and Cm did correlate in a manner that supports enhanced TC neuron excitability in D2 mice with elevated IOP.

At 9 months, there were no significant differences in Vrest, Rin, or Cm between D2 and D2-controls. For instance, Vrest was −77.7 + 1 mV for 9m D2 and −79.7 + 0.6 mV for D2-controls (p=0.25). Cm was notably lower in D2s (86.7 + 5.4 pF) compared to D2-controls (100.8 + 3.9 pF), but the difference was not significant (p=0.27). Likewise, Rin appeared elevated in D2’s (333 + 23 MΩ) compared to D2-controls (262 + 17 MΩ), but not significantly so (p=0.22).

As for AUC and I_50_, above, we noted that there was considerable variability in Vrest, Rin, and Cm. Therefore, we tested whether there was a correlation of each parameter with IOP in D2 mice (Figure 3F&G). For mean values of cells recorded from 9m D2 mice, Cm was negatively correlated with IOP (p = 0.025), while Rin was positively correlated with IOP (p = 0.011). In contrast, there was no significant correlation of Vrest with IOP (p = 0.96). Moreover, neither Cm, Rin, nor Vrest were significantly correlated with IOP at 4 months (p = 0.31, p = 0.78, p = 0.58, respectively). Thus, in 9m D2 mice, there was an association of several TC neuron parameters with the cumulative IOP such that cells from mice with higher IOP were more excitable, had a higher Rin, and had a lower Cm.

### TC neuron soma size is reduced in aged D2 mice

A reduction in cell size can contribute to increased Rin and would be consistent with the detected decrease in Cm. We next sought to verify this electrophysiological finding using a parallel anatomical approach in which we identified neuronal cell bodies in the dLGN using a NeuN antibody and measured their cross-sectional area in D2 and D2-control tissue at 4m and 9m (Figure 4). GABAergic interneurons were identified by staining with a GAD65/67 antibody and were excluded in order to focus this analysis on the excitatory TC relay neurons in the dLGN. At 4m, there was small shift in the median TC neuron soma area, from 229 μm^2^ in D2 control (interquartile range, IQR: 189-274 μm^2^; n = 214 cells from 6 mice) to 196 μm^2^ in D2 mice (IQR: 170-234 μm^2^; n = 106 cells from 4 mice; p = 0.00055, K-S test). At 9m, the difference was much more pronounced, with the median soma area being 233 μm^2^ in D2-controls (IQR: 196-280 μm^2^; n = 312 cells from 10 mice) and the median soma area being 167 μm^2^ in D2 mice (IQR: 135-199 μm^2^; n = 321 cells from 9 mice; p<0.00001, K-S test). This anatomical approach supports electrophysiological findings that TC neuron size is reduced in the 9m D2 dLGN and suggests that excitability changes at 9m, but not 4m, might be linked to TC neuron atrophy.

**Figure 4.**
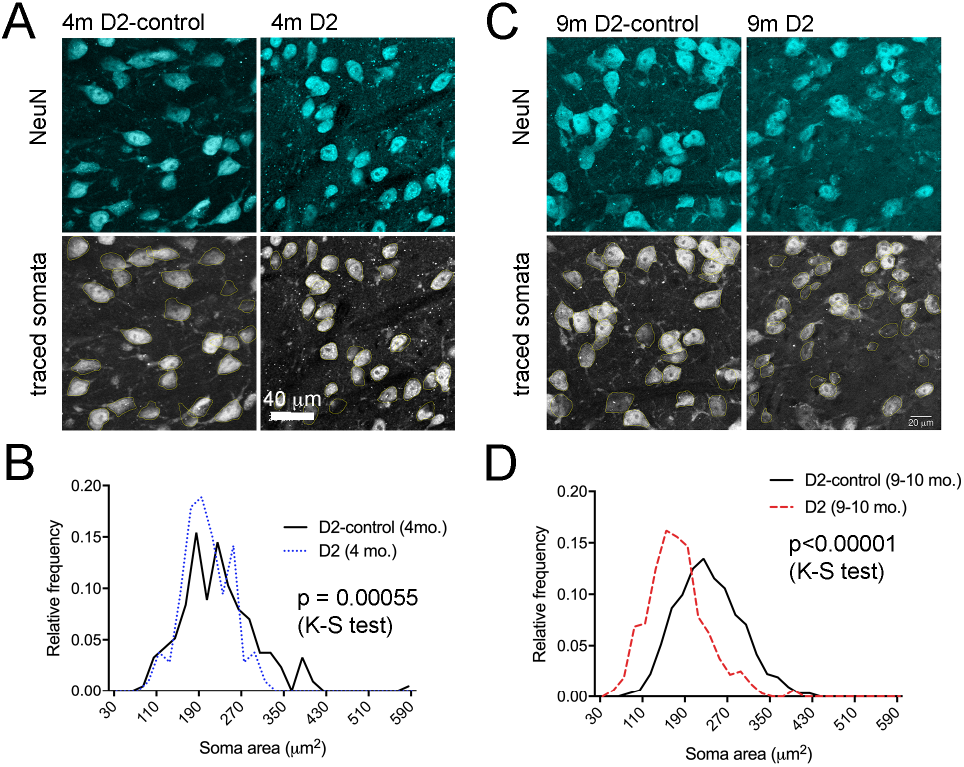
Measurement of TC neuron soma size in 4m and 9m D2 mice. 50 micron dLGN sections were stained using a NeuN primary antibody to label neurons while a GAD65/67 antibody was used to identify and exclude GABAergic interneurons. A&C) After imaging, TC neuron somata were traced. Lower panels show the somatic outlines (yellow). B) At 4m, TC neurons were slightly smaller in D2 mice (n = 106 cells from 4 mice, compared to D2-controls (n = 214 cells from 6 mice), although the difference did not pass our significance threshold for the K-S test (p = 0.00055). At 9m, there was a significant leftward shift in the distribution of TC neuron soma size in D2 mice (n = 321 cells from 9 mice) compared to D2-controls (n = 312 cells from 10 mice; p<0.00001, K-S test).

### Enhancement of TC Neuron excitability in an inducible ocular hypertension (OHT) model

To test whether the increase in dLGN TC neuron excitability is a common feature of experimental glaucoma or instead a quirk of the D2 mouse, we next turned to an inducible model in which eye pressure is elevated by injection of microbeads into the anterior chamber (Sappington et al., 2010; Calkins et al., 2018; Bhandari et al., 2019). This led to an approximately 5 mmHg increase in IOP over baseline (~30% increase; Figure 5A), which we have shown previously to trigger changes in the function of retinogeniculate (RG) synapses and TC neuron dendritic structure with minimal RGC loss five weeks after injection (Bhandari et al., 2019).

**Figure 5.**
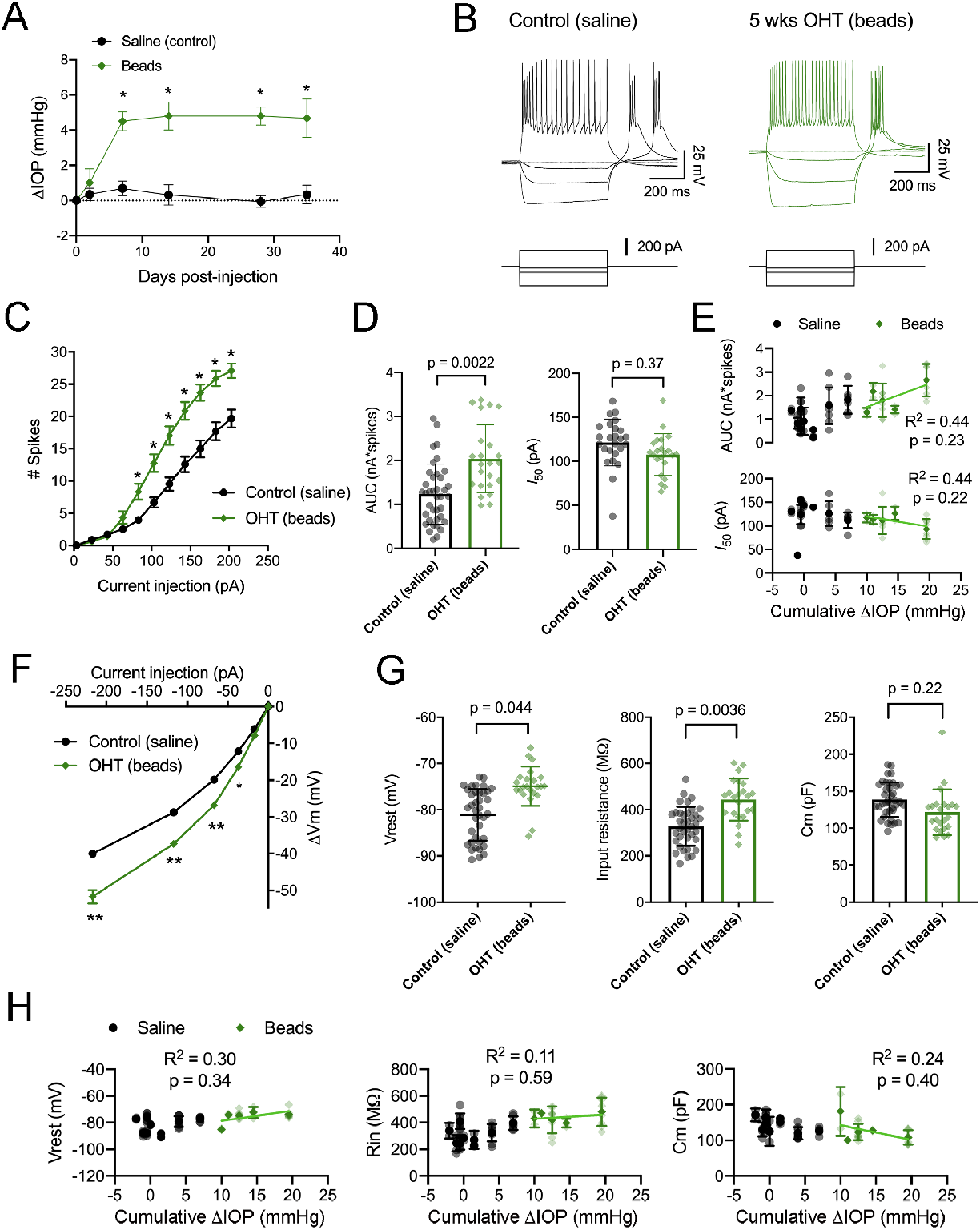
IOP elevation in the microbead model enhances dLGN TC neuron excitability. A) Following bilateral anterior chamber injection of 10-micron polystyrene microbeads, IOP was elevated by ~5 mmHg (n = 10 eyes from 5 mice) compared to saline-injected control mice (n = 18 eyes from 9 mice; *p<0.05, unpaired t-test). B) Whole-cell current clamp recordings from TC neurons in dLGN coronal slices show spiking and hyperpolarizing voltage responses to depolarizing (+180 pA) and hyperpolarizing (−20, −70, and −220 pA) current injections in TC neurons from saline-and bead-injected mice. C) Plot of mean + SEM of current-evoked spiking (*p<0.05, unpaired t-test). Saline data are from 38 cells from 9 mice while microbead data are from 23 cells from 5 mice. D) AUC was significantly elevated in cells from bead-injected mice compared to saline-injected controls (nested t-test). I_50_ was slightly reduced, although not significantly. Bars and errors bars show mean + SD while data points are measurements from individual cells. E) There was no significant correlation of AUC or I_50_ with IOP from bead-injected mice as assessed with a linear regression. Cumulative ΔIOP represents the between-eyes average of IOP measurements taken at 1 week, 2 weeks, and 4 weeks post-injection. F) Voltage responses to hyperpolarizing current injections were used to measure passive membrane properties from the same population of cells. G) TC neurons from bead-injected mice were slightly depolarized relative to saline-injected controls (nested t-test). Rin was significantly elevated and Cm was not significantly different. H) Passive membrane properties did not significantly correlate with IOP as assessed with a linear regression.

Here, we performed current clamp recordings to measure neuronal excitability and membrane properties from mice with microbead-induced OHT (Figure 5B), similar to experiments with D2 mice, above. One difference was that experiments with dLGN slices from bead-injected mice were recorded at room temperature rather than warmed, which changes the evoked spiking and membrane properties of TC neurons (Van Hook, 2020). Still, in bead-injected mice, we found that TC neuron excitability was enhanced when compared to TC neurons from saline-injected controls (Figure 5C-E). This was reflected as an increase in the number of action potentials evoked by depolarizing current injection (20-220 pA) and further demonstrated as an increase in the AUC, from 1.2 + 0.1 nA*spikes (n = 39 cells, 9 mice) to 2.0 + 0.2 nA*spikes (n = 24 cells, 4 mice; p=0.022). Although there was a trend toward a leftward shift in I_50_, (saline: 121 + 5 pA; beads: 108 + 5 pA), the difference was not statistically significant (p=0.37). Additionally, and similar to the 4m D2 mice, there was no significant correlation of either AUC (p = 0.23) or I_50_ (p =0.22) with IOP for bead-injected mice.

We next compared membrane properties, including Vrest, Rin, and Cm, between TC neurons from bead- and saline-injected mice as for D2 and D2-control mice, above (Figure 5F-H). We found that Vrest was slightly depolarized in TC neurons from bead-injected mice when compared to controls (beads: −74.9 + 0.6 mV, n = 24 cells, 5 mice; saline: −80.9 + 0.8 mV, n=39 cells, 9 mice; p=0.044). Additionally, Rin was elevated in bead-injected mice relative to saline-injected controls (beads: 443 + 20 MW; saline: 328 + 14 MΩ; p=0.0036). There was a trend toward a reduced Cm (beads: 121 + 4 pF; saline: 138 + 5 pF), but the difference was not significant (p = 0.22). Also, similar to the 4m D2 mice, there was no significant correlation of Vrest (p=0.34), Rin (p=0.59), or Cm (p=0.40) with IOP in microbead-injected mice.

### Reduced mEPSC frequency in D2 mice

We have previously shown that IOP elevation in the microbead model leads to a reduction in the frequency of miniature excitatory post-synaptic currents (mEPSCs) recorded from dLGN TC neurons (Bhandari et al., 2019). Therefore, we next used whole-cell voltage-clamp recordings of dLGN TC neurons from 4m and 9m D2 mice to probe whether and how excitatory inputs are altered in these mice as well (Figure 6). For 4m mice (Figure 6A-C), we found that mEPSC amplitude was similar in D2 and D2-control mice (D2-control: 8.3 + 0.6 pA, n = 12 cells, 5 mice; D2: 9.2 + 0.7 pA, n = 11 cells, 4 mice; p = 0.52). mEPSCs arising from cortical inputs are slower than those from RGC inputs due to their concentration at more distal dendritic sites and consequent dendritic filtering (Williams and Mitchell, 2008). Shifts in relative proportions of cortical-vs-retinal inputs can be reflected in changes in the mEPSC decay kinetics as occurs in monocular deprivation (Krahe and Guido, 2011). However, we did not observe any significant shift in mEPSC decay time constant in 4m TC neurons (D2-control: 2.1 + 0.2; D2: 2.3 + 0.3 ms; p = 0.74). The mEPSC frequency at 4 months was slightly reduced (D2-control: 19.7 + 2.7 Hz; D2: 13.0 + 2.3 Hz). Although the difference was not significant when assessed with a nested t-test (p = 0.22), the distribution of inter-event intervals was significantly different between D2-control and D2 (p<0.00001, K-S test).

**Figure 6.**
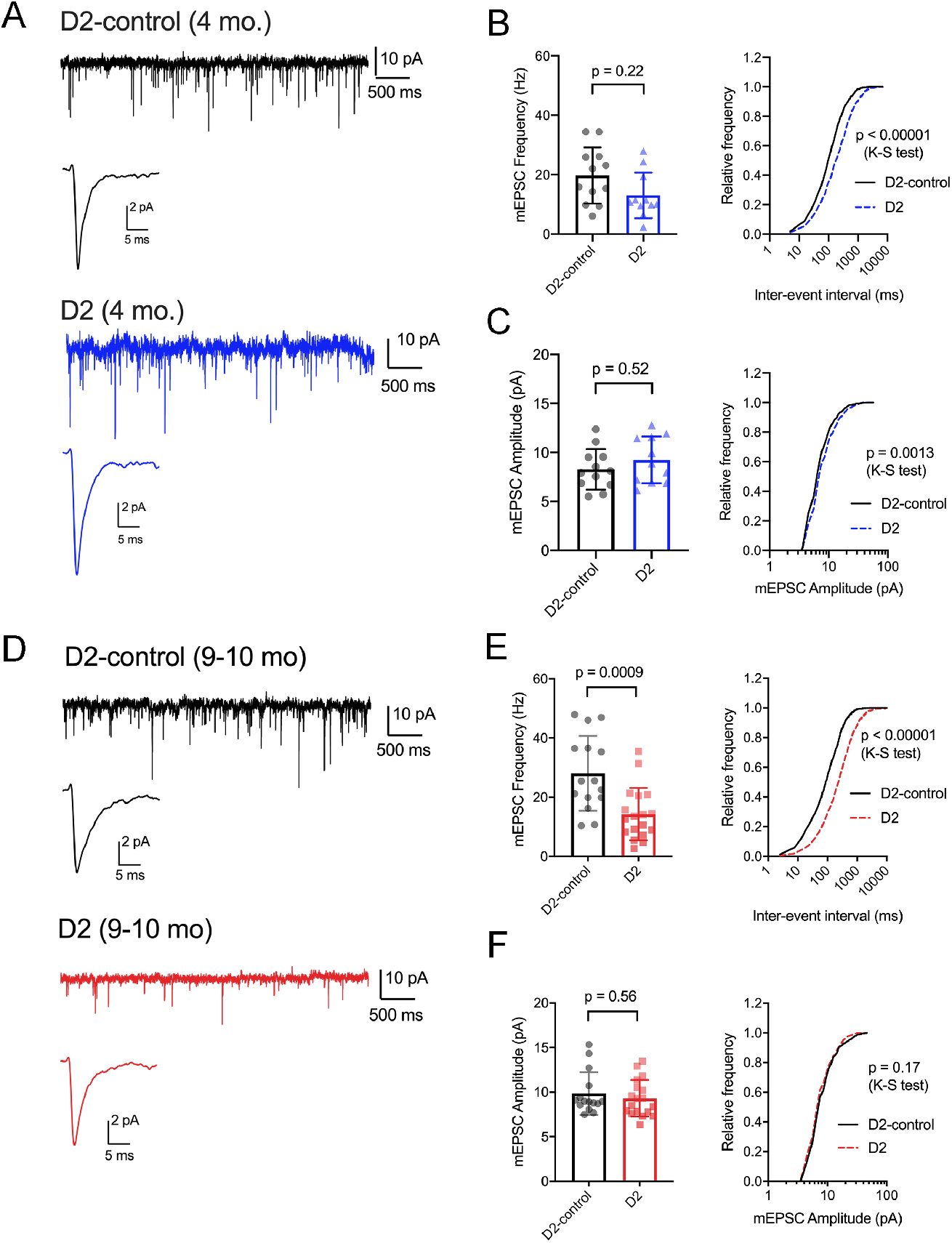
mEPSC frequency was reduced in D2 mice. A) Whole-cell voltage clamp records of mEPSCs recorded in the absence of stimulation for 4m D2-control (n = 12 cells from 5 mice) and D2 mice (n = 11 cells from 4 mice). Single events are the average of events detected in each cell. B) Mean + SD of mEPSC frequency was slightly reduced in 4m D2 mice. Individual points show the average instantaneous frequency for all events from individual cells. The difference was not statistically significant when assessed with a nested t-test, although there was a slight leftward shift in the distribution of inter-event intervals that was significant when assessed with a K-S test. C) mESPC amplitude was not different between 4m D2-controls and D2 mice. D) mEPSCs recorded from TC neurons from 9m D2-control (n = 15 cells from 4 mice) and D2 mice (n = 18 cells from 6 mice). E) mEPSC frequency was reduced at 9m. F) mEPSC amplitude was similar in D2-control and D2 TC neurons at 9m.

At 9m (Figure 6 D-F), mEPSC amplitude was also similar between D2 and D2-control mice (D2-control: 9.8 + 0.6 pA, n = 15 cells, 4 mice; D2: 9.3 + 0.5 pA, n = 18 cells, 6 mice; p = 0.56). Additionally, the mEPSC decay time constants were not significantly different at 9m (D2-control: 1.98 + 0.09 ms; D2: 1.86 + 0.11 ms; p = 0.84). mEPSC frequency was significantly lower in 9m D2 mice compared to D2-controls (D2-control: 28 + 3.3 Hz; D2: 14.3 + 2.1 Hz; p = 0.0009). This was also reflected as a significant shift in the cumulative distributions of inter-event intervals (p<0.00001, K-S test). In contrast to our findings with excitability and membrane properties, there was no significant correlation of mEPSC frequency with the three-month cumulative IOP (R^2^ = 0.48; p = 0.13).

### Loss of RGC axon terminals in the dLGN of D2 mice

Prior studies have indicated that RGC axon terminals in the superior colliculus (SC) are lost fairly late in disease in DBA/2J and microbead-injected mice and that terminal loss is preceded by swelling followed by atrophy (Crish et al., 2010; Smith et al., 2016). Additionally, we have shown that five weeks of a relatively modest and sustained OHT triggered by anterior chamber microbead injections does not have any effect on RGC axon terminal size or density in the dLGN (Bhandari et al., 2019).

Next, we sought to determine whether the size or density of RGC axon terminals, stained with an antibody raised against vGlut2, which is a selective label for RGC axon terminals in the dLGN, were altered in D2 mice at young age (4m) and in older mice (9m) with elevated eye pressure (Figure 7). In sections from 4m D2 mice (Figure 7A&B), we found that vGlut2 punctum density was similar to D2-control mice (D2: 11.2 + 0.8 puncta/1000 μm^2^, n = 4 mice; D2-control: 11.9 + 0.2 puncta/1000 μm^2^, n = 6 mice; p = 0.48). We found no evidence for RGC axon terminal atrophy or swelling, as punctum size was similar in controls and D2 (D2: 19.0 + 0.4 μm^2^; D2-control; 18.3 + 0.2 μm^2^, p = 0.21). Further supporting this, the cumulative distribution of vGlut2 punctum size was similar in 4m D2 and D2-control mice (p=0.18, K-S test).

**Figure 7.**
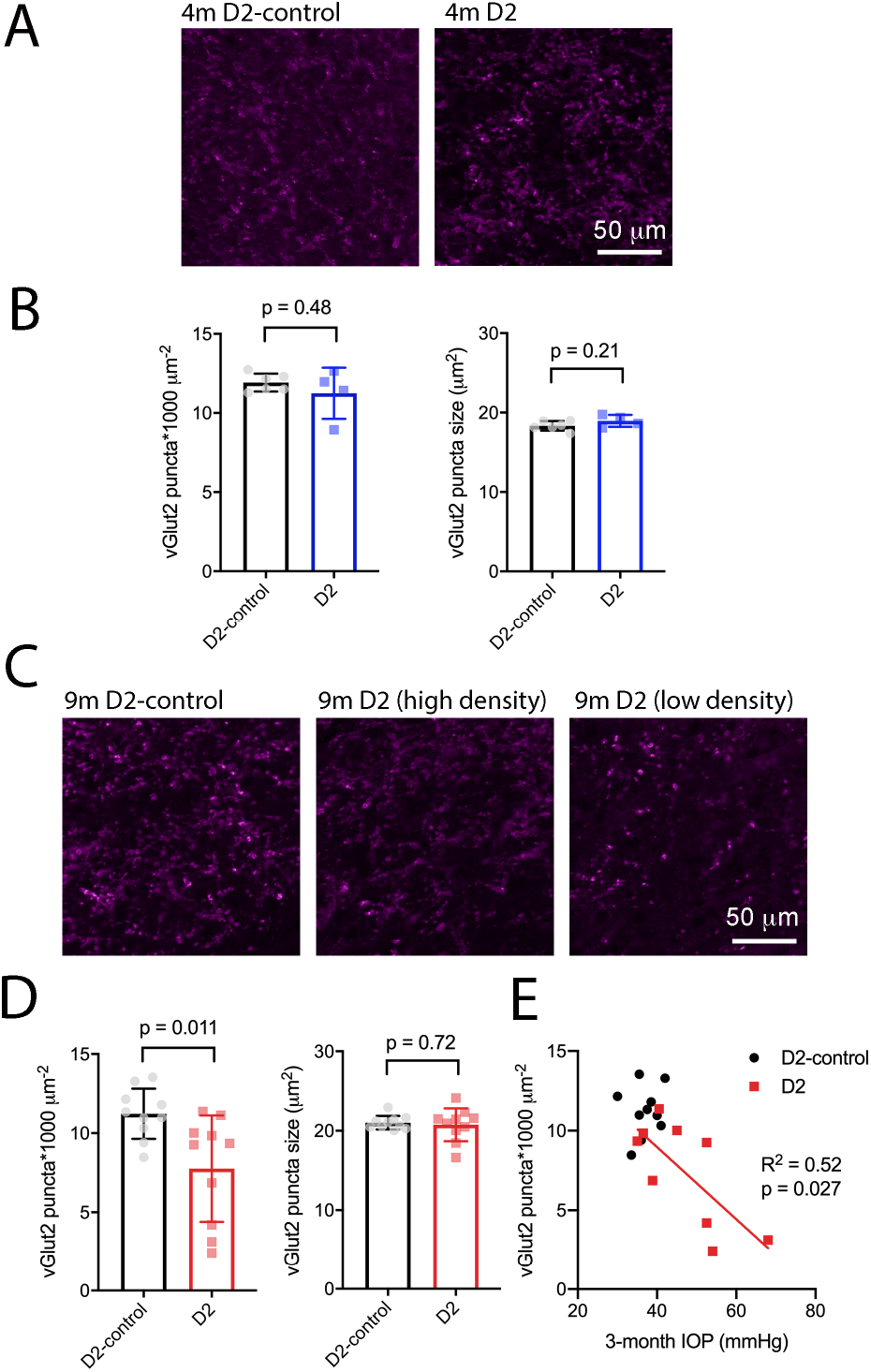
vGlut2-labeled RGC axon terminals are lost in an IOP-dependent manner in 9m D2 mice. A) 2.5-micron-thick maximum intensity projections of vGlut2 staining from the dLGN core of 4m D2-control (n = 6 mice) and D2 mice (n = 4 mice). B) Neither density nor size of detected vGlut2 puncta was significantly different between D2-control and D2 mice at 4m. C) vGlut2 images from 9m D2-control (n = 10 mice) and D2 mice (n = 10 mice). For D2 mice, images are shown from mice with higher vGlut2 density (middle panel) and lower vGlut2 density (right panel) to illustrate the range of labeling seen. D) vGlut2 density is significantly reduced in 9m D2 mice compared to D2-controls (unpaired t-test) while vGlut2 puncta size is not different. E) vGlut2 density in 9m D2 mice dLGN sections significantly correlated with the three-month cumulative IOP as assessed with a linear regression.

Similar to results at 4 months of age, there was no change in vGlut2 punctum size at 9m (D2: 20.8 + 0.44 μm^2^, n = 10; D2-control: 21 + 0.3 μm^2^, n = 10; p = 0.72; Figure 7C&D). Likewise, the cumulative distribution of vGlut2 punctum size was similar (p = 0.82, K-S test). Although punctum size was unchanged, vGlut2 density was significantly reduced in D2 mice compared to controls (Figure 7D; D2: 7.8 + 1.2 puncta/1000 μm^2^; D2-control: 11.2 + 0.5 puncta/1000 μm^2^; p = 0.011). Notably, there was considerable variability in vGlut2 punctum density, with a range of 2.4-11.4 puncta/1000 μm^2^ in D2 compared to 8.5-13.5 puncta/1000 μm^2^ in controls. To test whether the diversity in vGlut2 density in D2 mice might be related to IOP, we tested for correlation of vGlut2 punctum density with the three-month cumulative IOP (Figure 7E). Indeed, vGlut2 density was negatively correlated with the three-month cumulative IOP (p = 0.027), suggesting that a greater loss of RGC axon terminals is associated with a higher cumulative IOP in 9m D2 mice.

### Eye-pressure-associated RGC loss in D2 mice

Although early changes to the structure and function of RGCs, their axons, and their projection targets in the brain are likely to contribute to visual impairment in glaucoma, RGC degeneration is thought to be the major cause of irreversible vision loss. Therefore, we sought to determine the differences in RGC degeneration in D2 mice at 4m and 9m of age and relate RGC degeneration to IOP in order to have a marker of glaucoma severity as it relates to IOP, mouse age, and the functional changes to synapses and TC neuron spiking behavior in the dLGN. To accomplish this, we labeled RGCs by immunofluorescence staining for RBPMS, a selective RGC marker and counted RGCs to determine their density in four quadrants of the central and peripheral retina (Figure 8).

**Figure 8.**
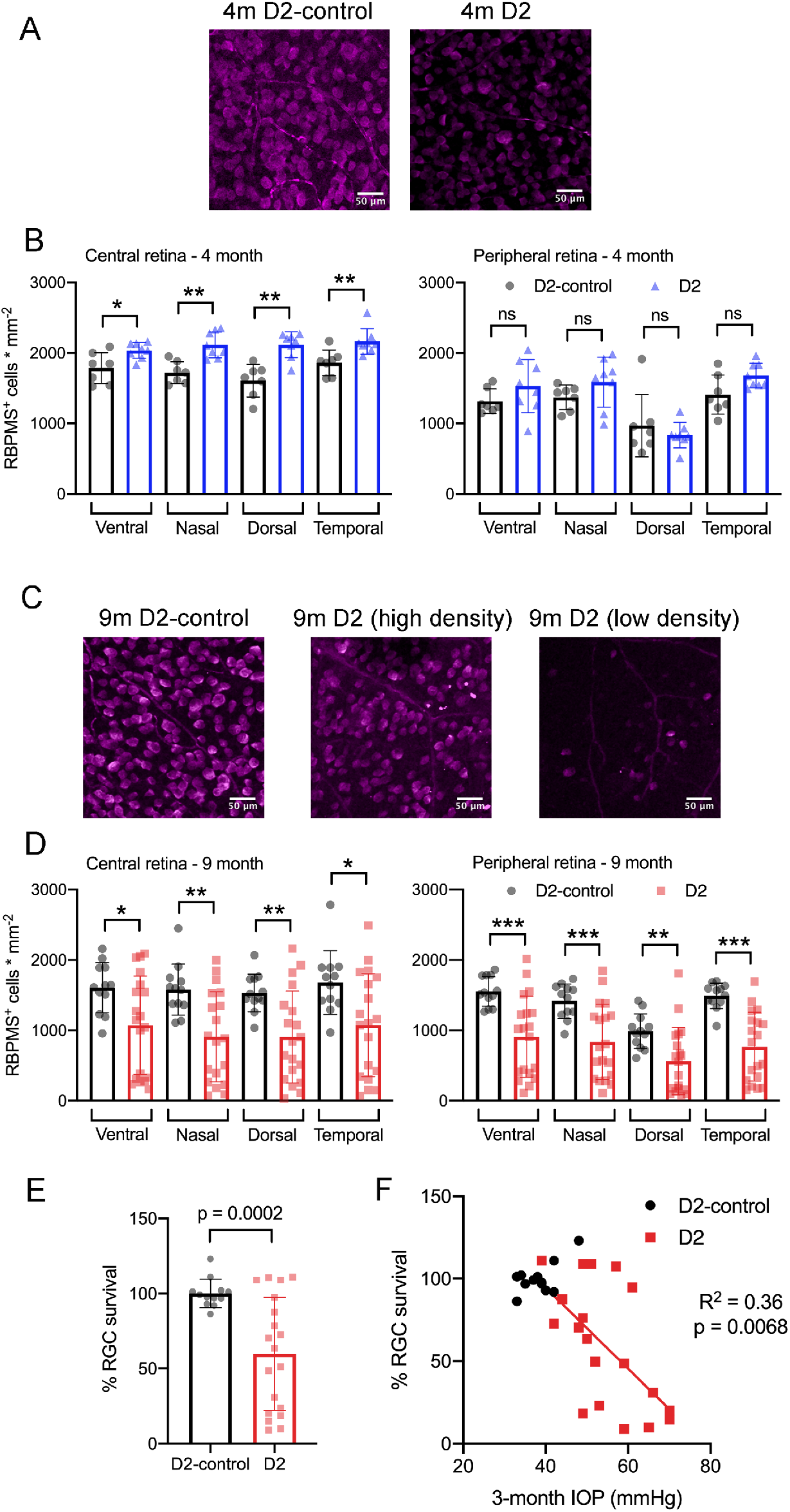
RBPMS-stained retinal ganglion cells are lost in an IOP-dependent manner in 9m D2 mice. A) Immunofluorescence images from RBPMS-stained retinas from a 4m D2-control and a D2 mouse show similar RBPMS+ RGC soma density. B) Quantification of RBPMS+ cell bodies in four quadrants of central and peripheral retina. Density was slightly higher in 4m D2 mice (8 retinas from 4 mice) compared to D2-control mice (n = 7 retinas from 4 mice) in central retina (*p<0.05, unpaired t-test). Bars and error bars represent mean + SD while individual points are RGC density measurements from individual retinas. C) RBPMS images from 9m D2-control (left) and D2 mouse retinas (center and right). Images from 9m D2 retinas with higher and lower RBPMS+ cell density are shown to illustrate the diversity of RGC densities seen in D2 mice. D) RBPMS+ cell density was lower in all four quadrants of central and peripheral retinas of 9m D2 mice (n = 19 retinas from 12 mice) compared to D2-controls (n = 12 retinas from 7 mice) illustrates the diversity of RGC survival from mouse-to-mouse. Bars and error bars represent mean + SD while individual data points are RGC survival in individual retinas. E) Quantification of RGC survival across the entire retina at 9m. F) RGC survival significantly correlated with the three-month cumulative IOP in 9m D2 mice, as assessed using a linear regression.

At 4m (Figure 8A&B), we found that there was no significant reduction in RGC density in D2 mice compared to D2-controls in either central or peripheral retina (D2: n = 8 retinas; D2-control: n = 7 retinas). Rather, in the central retina, the RGC density in D2 mice was slightly higher than RGC density in D2-controls. In peripheral retina, however, there was no significant difference between D2- and D2-control RGC density in any of the four quadrants. At 9m, however, RGC density was significantly lower in D2 than D2-controls (D2: n = 19 retinas; D2-control n = 12 retinas). This was the case in both central retina and peripheral retina.

Across the retina, there was more variability in the RGC density in D2 mice compared to D2-controls at 9m (Figure 8C-F). For instance, the standard deviation of RGC densities across four quadrants in central and peripheral retina in D2-control mice was 289 cells/mm^2^ while it was 599 cells/mm^2^ for age-matched D2-mice. We then tested whether the differences in RGC density were associated with IOP by calculating RGC survival as % of control RGC density and testing for correlation of RGC survival across the entire retina with the three-month cumulative IOP. Indeed, RGC survival was negatively correlated with 3-month cumulative IOP (p = 0.0068), supporting a relationship between eye pressure and RGC loss in 9m D2 mice.

## Discussion

In this study, we identify functional changes to the TC relay neurons and their synaptic inputs in the dLGN of DBA/2J mice, a widely-used and established rodent model of inherited glaucoma. Specifically, we find in younger D2 mice that do not have any detectable OHT or RGC loss, TC neurons more readily fire action potentials in response to depolarizing current stimuli and that this is associated with a slight depolarization and increase in neuronal input resistance. TC neuron excitability and membrane parameters were only subtly different from controls in 9-month D2 mice. There was much more variability in neuronal excitability and passive membrane properties at 9m and these parameters correlated with D2 mouse IOP. In mice with a modest OHT triggered by anterior chamber microbead injection, TC neuron excitability was also increased in a manner that resembled the 4-month D2 mouse population in that it was associated with a small depolarization and increase in Rin without significant change in Cm. Moreover, excitability and passive membrane properties (AUC, I_50_, Cm, Rin) did not correlate with IOP in microbead-injected mice, similar to 4m D2 mice.

With the differences in functional effects observed in 4m and 9m D2 mice as well as in consideration of the effects in microbead-injected mice, we suggest that the pattern of functional changes in the dLGN represents distinct, although potentially overlapping, phases of glaucoma’s effects. At 4m, the effects are likely to represent homeostatic plasticity (Turrigiano, 2011, 2012; Crish and Calkins, 2015); there was no detectable RGC loss at this time point and IOP remained low, similar to control mice. However, other studies have documented that functional changes can occur in the visual pathway of D2 mice at this age and earlier including diminishment of optic nerve transport, remodeling of astrocytes in the optic nerve, and tau hyperphosphorylation (Crish et al., 2010; Dengler-Crish et al., 2014; Crish and Calkins, 2015; Cooper et al., 2016; Wilson et al., 2016). Thus, the enhanced excitability in 4m D2 mice might represent functional homeostasis triggered as a downstream consequence of compromised optic nerve health and disrupted signal transmission from RGCs.

Indeed, homeostatic upregulation of intrinsic excitability has been documented in response to altered visual input in visual cortex (Maffei and Turrigiano, 2008; Lambo and Turrigiano, 2013). OHT also alters RGC spike output, causing an early increase in excitability due to changed Na^+^ channel expression (Risner et al., 2018, 2020; McGrady et al., 2020). Other studies have documented diminishment of spontaneous and light-evoked RGC spike output occurring at different times with different IOP manipulations (Della Santina et al., 2013; Ou et al., 2016; Risner et al., 2018; Bhandari et al., 2019). Changes in RGC spike output might alter depolarization-triggered CREB activity in TC neurons (Pham et al., 2001; Guido, 2008; Krahe et al., 2012; Dilger et al., 2015). Altering RGC spiking might also influence activity-dependent release of the neurotrophin BDNF, which is important in glaucoma pathology (Crish et al., 2013; Domenici et al., 2014; Gupta et al., 2014; Dekeyster et al., 2015; Valiente-Soriano et al., 2015), regulates development of retinal ganglion cells and their brain projections (Marshak et al., 2007; Cohen-Cory et al., 2010; Nikolakopoulou et al., 2010), and is involved in homeostatic and experience-dependent plasticity of cortical neurons (Rutherford et al., 1998, 1998; Desai et al., 1999; Bracken and Turrigiano, 2009).

In contrast to our findings at 4m, there was a notable correlation of neuronal excitability, including spiking parameters and passive membrane properties with IOP in 9m mice. In these cases, higher IOP was associated with increased TC neuron excitability in the dLGN. Moreover, RGC density was significantly reduced across the retina by 9m and we found that higher IOP was also associated with lower RGC survival. At this time point, we observed electrophysiological signatures of reduced TC neuron size and confirmed the TC neuron atrophy by measurements of TC neuron soma area. This is consistent with other studies of retinal projection targets in fairly late-stage glaucoma from human patients and primate and rodent animal studies which have also documented neuronal atrophy in the dLGN and SC (Yücel et al., 2001; Gupta et al., 2007, 2009). Thus, the pattern of functional changes we observed might represent a shift from pre-degenerative homeostasis to pathological dysfunction associated with RGC somatic and axon terminal degeneration (Orr et al., 2020).

Probing synaptic function, we also found in both younger and older D2 mice that the excitatory synaptic input onto dLGN TC neurons was reduced, as indicated by a reduction in mEPSC frequency. There was no change in mEPSC amplitude, suggesting that the post-synaptic AMPA receptor population was stable at excitatory synapses onto TC neurons. Moreover, in older mice, there was a reduction in the density of vGlut2-labeled RGC axon terminals in the dLGN in a manner that correlated with IOP such that higher eye pressure was associated with a greater terminal loss.

Previous studies of the superior colliculus of D2 mice have reported that RGC axon terminals survive until fairly late in the disease and only degenerate well after RGC loss (Crish et al., 2010; Smith et al., 2016; Wilson et al., 2016). In our sample of 9m D2 mice, we found that RGC axon terminals were lost in an IOP-dependent manner. We did not see evidence of either RGC axon terminal atrophy or swelling, as has been reported in ultrastructural studies of the SC (Smith et al., 2016). Several possibilities could account for this difference. First, it is possible is that, unlike in the SC, RGC terminals in the dLGN do not exhibit phases of swelling and atrophy prior to degeneration. This could represent a difference between these two visual structures, possibly resulting from relative proximity of each structure to the retina, which is known to be associated with differential effects on the optic projection (Crish et al., 2010; Calkins, 2012). Alternatively, light microscopy as we implemented it might be too coarse of an imaging modality to detect subtle changes in RGC axon terminal structure if OHT triggers early swelling or later atrophy. Future ultrastructural studies of the dLGN would be able to address this possibility. Notably, Smith and colleagues combined their measurements of axon terminal size with anterograde transport studies and found that areas of the SC with intact transport from the retina tended to have terminal swelling while areas deficient in transport (taken as a sign of poorer health) tended to have atrophied terminals (Smith et al., 2016). We did not perform transport experiments as part of this study. Doing so in combination with vGlut2 staining in future studies might reveal a similar pattern in the dLGN.

Although retinogeniculate (RG) synapses provide the major excitatory drive for TC neurons, they comprise only ~10% of their total number of excitatory synaptic inputs (Bickford et al., 2010; Guido, 2018). The majority of excitatory inputs onto TC neurons are the result of corticothalamic (CT) synapses arising from layer VI of the visual cortex. These synapses function to modulate TC neuron responsiveness (Sherman and Guillery, 2002, 2011) and have a low release probability, as evidenced by their notable synaptic facilitation occurring during repeated stimulation (Turner and Salt, 1998; Jurgens et al., 2012). We find here that TC neurons from D2-control mice have a baseline mEPSC frequency of approximately 25 Hz, which is similar to what we have shown previously in recordings at ~30-33°C (Van Hook, 2020), although higher than mEPSC frequency detected in recordings at room temperature (Bhandari et al., 2019; Van Hook, 2020). In the current study, we found that mEPSC frequency recorded in TC neurons was reduced in both 4m and 9m D2 mice compared to their age-matched controls. The mechanisms underlying the reduced mEPSC frequency are unclear. IOP was not elevated in our sample of 4m D2 mice, nor was mEPSC frequency correlated with IOP in 9m D2 mice, in contrast to electrophysiological markers of TC neuron excitability, vGlut2 loss, and RGC loss. RGC axonal transport, cytoskeleton, and energy homeostasis is disrupted at fairly young ages in D2 mice, and this is accompanied by morphological changes in RGC axon terminal mitochondria in the SC (Crish et al., 2010; Smith et al., 2016). Synaptic transmission requires ample energy supply and functioning mitochondria to handle vesicle recycling, regulation of presynaptic vesicle pool size and release probability, and presynaptic Ca^2+^ handling (Ly and Verstreken, 2006). Therefore, perturbations of RGC axon terminal mitochondria and axonal energy supply are likely to influence synaptic output and might contribute to our observed synaptic changes.

In a previous study using the microbead approach to induce OHT, we suggested that a similar reduction in mEPSC frequency might be attributable to a loss of post-synaptic TC neuron synapses, as evidenced by a reduction in TC neuron dendritic complexity (Bhandari et al., 2019). Dendritic atrophy in retinorecipient neurons in late-stage glaucoma has also been documented elsewhere, yet we have not yet tested whether or along what time course this occurs in D2 mice.

Alternatively, changes in mEPSC frequency might arise from changes to non-RGC inputs. Krahe & Guido have shown that monocular deprivation leads to a homeostatic increase in synaptic input arising from the CT pathway in the dLGN (Krahe and Guido, 2011). They reached this conclusion from observing changes in short-term plasticity during CT tract stimulation and an increase in the proportion of detected mEPSCs with slower decay kinetics, which results from filtering of signals arising at distal dendritic sites (Williams and Mitchell, 2008). It remains to be seen what proportion of TC neuron mEPSCs arise from retinal vs. cortical sources and whether OHT and optic nerve pathology can influence CT synaptic function. CT innervation is shaped by RG inputs during development (Seabrook et al., 2013) and dysfunction and injury to RGC axons might re-awaken those developmental processes in adulthood (Nahmani and Turrigiano, 2014). However, we did not detect any changes in mEPSC kinetics at either 4m or 9m, arguing against up- or down-regulation of CT input. Given the loss of vGlut2-labeled RGC axon terminals in 9m D2 mice, it is likely that some reduction in mEPSC frequency is the result of less RGC input to each TC neuron at that time point, but not in 4m mice, when vGlut2 labeling was unchanged compared to controls.

It will ultimately take examination of the properties of CT and RG synaptic function resulting from optic tract or CT tract stimulation to test for relative changes in synaptic function in D2 mice. We previously used optogenetic activation of RGC axons in microbead-injected mice to show that presynaptic vesicle release probability at RG synapses was increased in OHT (Bhandari et al., 2019). This was prior to substantial RGC degeneration or any detectable loss of vGlut2 staining in the dLGN, suggesting it might be an early, homeostatic response to pressure elevation. Numerous RGCs provide convergent synaptic input to each TC neuron, although only ~3 of those inputs are responsible the majority of excitatory drive to each cell (Chen and Regehr, 2000; Hammer et al., 2015; Litvina and Chen, 2017; Rompani et al., 2017). Given the IOP-associated decline in vGlut2 staining in 9m D2 mice, we do predict that later stages of disease will be characterized by a reduction in total RG synaptic strength, possibly as a reduction in RGC convergence to each TC neuron, although the strength of inputs from single fibers is likely to be diminished as well as axonal and synaptic function becomes more compromised.

In conclusion, prior studies have documented numerous changes to the structure of RGC axon terminals as well as to post-synaptic neurons in visual structures in the brain, especially in the SC, at various time points in glaucoma. The findings in the current study shed light on the progression of functional changes at the cellular and synaptic scale in the dLGN and relate those changes to established markers of glaucomatous progression – namely IOP and RGC loss in the retina. This study enhances the picture of disease progression and provides important information linking the IOP to vision loss in glaucoma.

## Acknowledgements

Elizabeth Bierlein and Ashish Bhandari for comments on the manuscript.

## Author Contributions

MJVH: Funding, experimental design, conducting experiments, analyzing data, writing manuscript

CM: conducting experiments, analyzing data

JCS: conducting experiments, analyzing data

